# Modeling Transitions between Responsive and Resistant States in Breast Cancer with Application to Therapy Optimization

**DOI:** 10.1101/031245

**Authors:** Chun Chen, John J. Tyson, William T. Baumann

## Abstract

We present a mathematical model that captures the transitions among three experimentally observed estrogen-sensitivity phenotypes in breast cancer cells. Based on this model, a population-level model is created and used to explore the optimization of a therapeutic protocol

## I. Introduction

Breast cancer is one of the most common cancers in women and approximately 70% of these cancers express the estrogen receptor (ERα). The growth of many of these cancers is driven by estrogen, which binds to ERα, a potent transcription factor, up-regulating proliferation. One form of endocrine therapy, aromatase inhibitors, reduces the biosynthesis of estrogen to shut down the estrogen receptor pathway and inhibit cell growth. Unfortunately, in many cases cells deprived of estrogen will eventually become able to grow under the deprived conditions, thus becoming resistant to the drug. Our interest is in understanding this transition to resistance and how to overcome it.

When cells grown in physiological levels of estrogen are transitioned to a medium with low estrogen, some of the cells will die while others become quiescent, ultimately adapt, and begin growing again. Similarly, if these growing cells are then transitioned to a medium with essentially no estrogen, some will die while others will ultimately begin to grow again. These results lead us to consider three estrogen sensitivity states: sensitive, hypersensitive, and independent. After hypothesizing a mechanism for these three sensitivity states, we built a simple stochastic mathematical model, based on an influence diagram drawn from the literature, to test the plausibility of our hypothesis. Suitable parameterization enabled the model to reproduce the qualitative behavior of cells with regard to estrogen sensitivity [1].

To examine the implications of this model for therapy, we derived a population model from our single-cell model. The population model accounts for the transitions of cells from one sensitivity state to another, as well as the growth and death rates of the cells in each state, as a function of the estrogen level. This high level model enabled us to easily examine the effects of aromatase inhibitor therapy on the population (tumor) and to optimize the parameters of the protocol to minimize overall growth.

## II. Results

The cell-level model has only two dynamic variables: active estrogen receptor at the membrane (ERM), and activated growth factor receptors (GFR). A potential landscape for the system is shown in Figure 1, where the dynamics of the system can be visualized as a point on the landscape that always moves in the downhill direction. The system has four basins of attraction where ERM^low^/GFR^low^ corresponds to the sensitive estrogen state, ERM^high^/GFR^low^ to the hypersensitive state, and GFR^high^ to the independent states.

**Figure 1:**
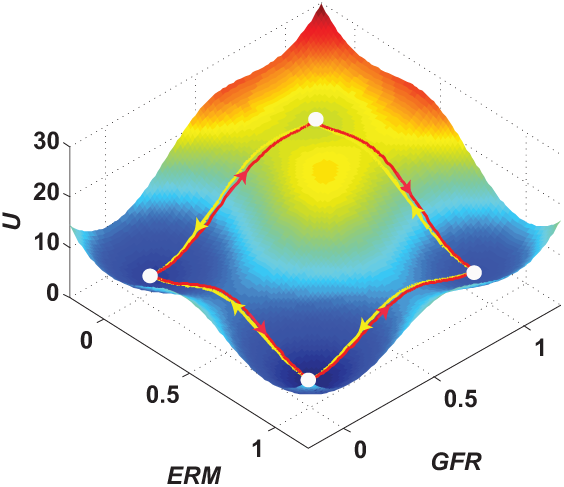
Potential landscape for a specific concentration of estrogen.

The population model allowed us to study the effect of changing estrogen levels on the numbers of cells in each basin. We cyclically deprived the cells of estrogen for a period and then restored estrogen for a period. The parameters of the protocol are the treatment time and the break time. Figure 2 shows the results of two different protocols. Optimization of the protocol parameters shows that case (b) is actually optimal in terms of minimizing population growth. In this case, less overall treatment than in case (a) leads to better results. With the parameters of our model we were not able to drive the number of cells to zero, but we could reach a state of no net growth.

This work provides a framework for using dynamic modeling for therapy optimization. Experimental quantification of the various transition, growth, and death rates will be necessary to determine what can realistically be

**Figure 2:**
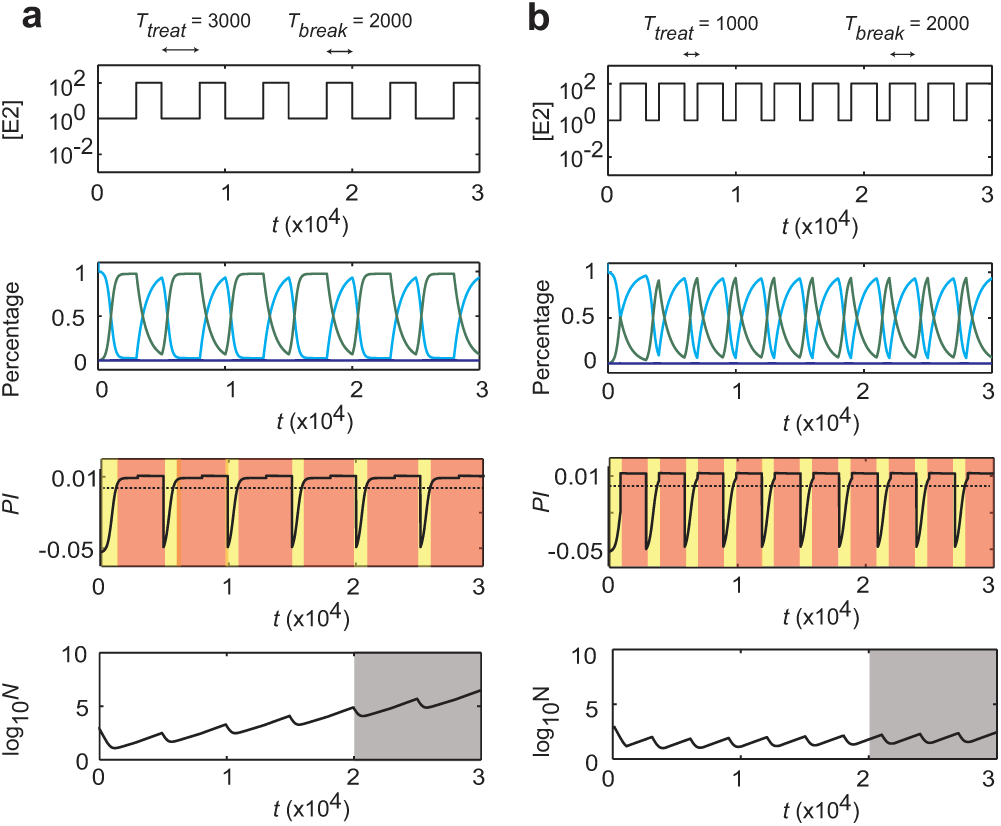
Results for two treatment protocols showing estrogen concentration (top), percentage of estrogen sensitive and hypersensitive cells, proliferation index of the population (>0 indicates growth), and the total number of cells in the population.

## III. Quick Guide to the Methods

The model describing the different estrogen sensitivity states is a phenomenological model, as the detailed mechanisms are unknown. The three components of the model are *E2ER,* estrogen-bound estrogen receptor, *ERM*, active membrane-associated estrogen receptor, and *GFR,* active growth factor receptors. The equations governing the model are given by:

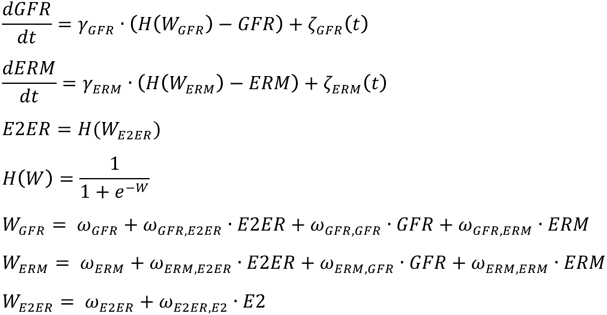

The model uses a simplified framework suitable for modeling systems at a phenomenological level. The framework, discussed in detail in [2], is based on soft-Heaviside functions, *H*, with arguments that are linear in the variables. The parameters of the model are the γ’s and ω’s. Since we are interested in stochastic transitions between steady states, we have added Gaussian white noise terms, ζ(t), to each equation. We assume the noise terms have the same autocorrelation, 〈*ζ*(*t*)*ζ*(*t*′)〉 = 2*D* · *δ*(*t* − *t*′), where *D* characterizes the magnitude of the random fluctuations.

To obtain an approximate potential landscape for visualizing our system we used ideas from [3]. Writing the above equations in vector form as

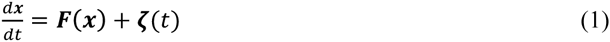
the Fokker-Planck equation for the probability density, *P(x,t),* is given by

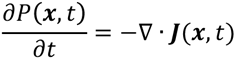
where J is the probability flux given by

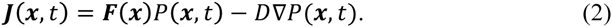

Solving for the steady state probability, where ∇ · **J** = 0, allows us to solve (2) for ***F*** as

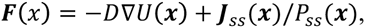
where *U*(*x*) = −ln (*P_ss_*(*x*)) is the approximate potential landscape of the system, and the dynamics of the system cause it to move in the direction of the negative gradient of the potential.

Similar to [4], we describe a population of cells by a vector, ***v***, of length *m* = 4. The *i*’th entry of the vector is the number of cells in the region *R_i_* surrounding the *i*’th steady state. The total number of cells in the population at time *t* is 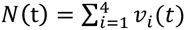. The population vector *v* evolves according to the equation *dv/dt* = *A · v* where *A* is an *m*×*m* transition rate matrix. The off-diagonal elements of *A*, *A_ij_* = *k_ij_,* are the transition rates from region *j* to region *i*, and the diagonal elements are *A_ii_* = *u_i_* − Σ*_j≠i_ k_ji_*, where *u_i_* is the proliferation rate of cells in region ***i***.

To estimate the transition rates *k_ij_* among the different phenotypic regions of the dynamical system, we followed the approach in [5] based on applying the Wentzell-Freidlin theory [6] to Eq. (1) for the ‘small noise’ case. The key idea of this theory is that the most probable transition path from *R_j_* at time 0 to *R_i_* at time *T,* 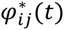 for *t* ∈ [0, T], minimizes the action functional

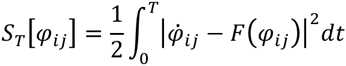
over all possible paths. This optimal path is referred to as the minimum action path (MAP). We are interested in MAPs between fixed-point attractors (stable steady states) in regions *R_1–4_.* The MAPs connecting each pair of attractors were computed numerically by applying the minimum action method used in [7].

The MAP 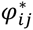 is useful for our purposes because we find that *k_ij_*, the transition rate from state *R_j_* to state *R_i_*, is related to 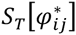 by the empirical equation

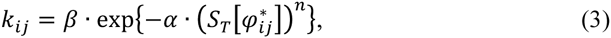
where the parameters *α, β* and *n* are estimated by fitting Eq. (3) to the results of direct Monte Carlo simulations of the dynamic model (1). After these parameters are fit, future calculations for the *k_ij_*, as the estrogen concentration changes, can be done deterministically by computing the MAP and using (3).

While these methods have been applied to a specific problem in cancer therapy, they have much wider applicability. The framework we used for modeling with ordinary differential equations has proven useful in many contexts, from modeling the cell cycle in yeast to examining the interaction between autophagy and apoptosis in mammalian cells. Also, the gradient landscape computation holds for any models of the form (1) of any dimension, but is probably most useful for visualizing models of dimension 2. Finally, the Wentzell-Freidlin theory can also be used for any models of the form (1).

